# Extensive single-cell genomics reveals bacterial diversity and diverse phage host ranges in the area in and around the Red Sea

**DOI:** 10.1101/2020.03.05.962001

**Authors:** Yohei Nishikawa, Masato Kogawa, Masahito Hosokawa, Ryota Wagatsuma, Katsuhiko Mineta, Kai Takahashi, Keigo Ide, Kei Yura, Hayedeh Behzad, Takashi Gojobori, Haruko Takeyama

## Abstract

To improve our understanding of the environmental microbiome, we developed a single-cell genome sequencing platform, named SAG-gel, which utilizes gel beads for single-cell isolation, cell lysis, and whole genome amplification (WGA) for sequencing. SAG-gel enables serial, parallel and independent reactions of > 100,000 single cells in a single tube, delivering high-quality genome recovery with storable randomized single-cell genome libraries. From soil and marine environmental sources, we acquired 734 partial genomes that are recapitulated in 231 species, 35% of which were assigned as high-to-medium qualities. We found that each genome to be almost unique and 98.7% of them were newly identified, implying the complex genetic diversities across 44 phyla. The various metabolic capabilities including virulence factors and biosynthetic gene clusters were found across the lineages at single-cell resolution. This technology will accelerate the accumulation of reference genomes of uncharacterized environmental microbes and provide us new insights for their roles.

## Introduction

Technological innovations in genome sequencing have improved our understanding of the genomic characteristics, genetic diversity, and metabolic capabilities of various biological samples. In particular, metagenomics and singlecell genomics exhibit great potential to enhance our understanding of the genomic structure and dynamics of complex microbial communities^1, 2^. The recent accumulation of metagenomic sequencing data and advances in computational techniques have enabled the integration of sequencing data, generating metagenome-assembled genomes (MAGs)^3, 4^. Although MAGs are expected to provide an expedient path for exploring microbial dark matter^5^, several problems remain including the inefficiency of genome reconstruction, 16S rRNA gene recovery^5^, and identification of microbial plasmids^6^.

To overcome these problems and confirm the reliability of sequencing data, single-cell genome sequencing plays an important role^7, 8^. Recently, microfluidic-based whole genome amplification (WGA) has been identified as a replacement for conventional fluorescence-activated cell sorting (FACS)-associated multi-well platforms^9, 10^, minimizing operating costs as well as the risk of DNA contamination, leading to increased reaction efficiency and throughput. In particular, droplet microfluidics is a noteworthy technique considered suitable for handling single cells^11–13^. However, the immiscible oil phase makes it difficult to conduct multistep reactions in droplets, causing an obstacle to lysis gram-positive and -negative bacteria and to exclude the inhibitors for WGA.

Herein we describe a method for multiplexed single-cell genome sequencing with massively produced single-cell amplified genomes (SAGs) in the gel that we named SAG-gel sequencing. We obtained high-coverage single-cell genome sequences of various environmental samples including soil, sediment, and seawater collected from the area surrounding the Red Sea in Saudi Arabia. This paper demonstrates that massively parallel single-cell genomics accumulates genome sequencing data of unknown species, unveils the obscured diversity present in uncultured environmental microbiome, and provides insight regarding their function in ecosystems.

## Results

### Workflow of SAG-gel platform

The strategy of SAG-gel is to perform a series of reactions on encapsulated single cells in a massively parallel manner (Fig. 1a). Single cells are massively captured in gel beads and lysed by enzyme cocktails designed for both gram-positive and - negative bacteria. Through the whole process, SAG-gel converts tiny genomes of various microbial cells into the amplified genomes in uniform-shaped beads floated in a single-tube. The gel matrix facilitates maintaining the compartmented genomes during cell lysis, washing, and WGA processes and preventing cross-contamination, and protecting the amplified DNA for long-term storage.

**Figure 1.**
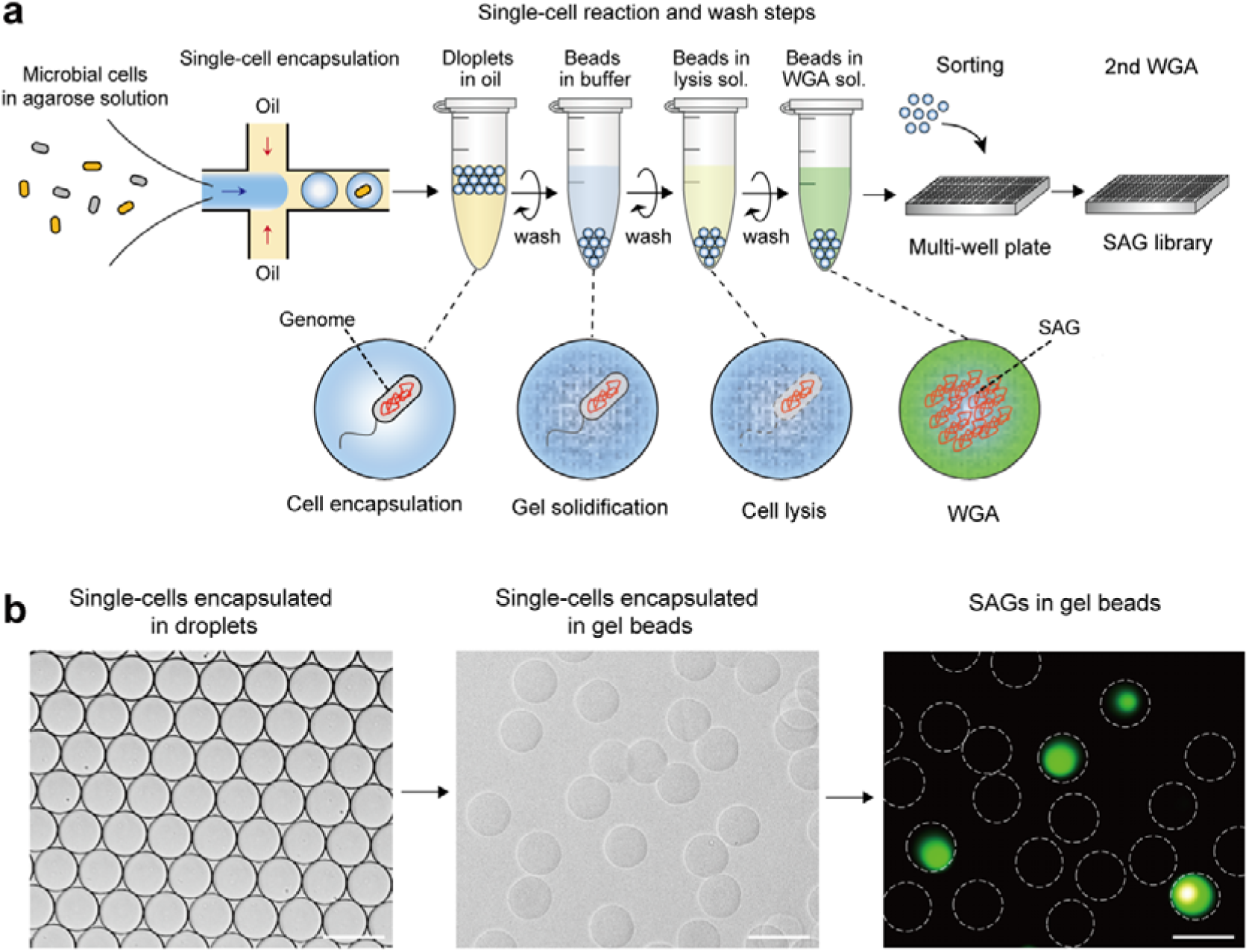
Single-cell amplified genome in the gel (SAG-gel) approach for massively parallel single-cell genome sequencing. a: Bacterial suspensions are encapsulated into 40 μm of microfluidic droplets at the single-cell level (0.1 cell/droplet) with ultra-low melting temperature agarose solutions. Collected droplets are solidified by cooling and single-cells are captured in gel matrix. Gel-beads are transitioned in aqueous phase, which enables repeating “reaction and wash” steps. Through these steps, single-cells were lysed and whole genome was amplified. Gel-beads after WGA can be isolated into multi-well plate with FACS, which can be applied to subsequent analysis including the next generation sequencing. b: Microscopic image of microfluidic water in oil (W/O) droplets and gel-beads. Amplified DNA can be visualized by adding DNA intercalating dye (SYBR Green). The scale bar is 50 μm.

Based on this concept, we developed a gel-beads based single-cell processing method by using gram-positive and -negative model bacteria (*Escherichia coli* and *Bacillus subtilis*). After cell encapsulation, gel beads were dispersed into the aqueous phase, which enables small molecules to penetrate into gel beads. By repeating a series of “reaction and wash” steps, practical step-by-step reactions including cell lysis, DNA purification, and WGA can be achieved (Fig. 1b). The gel beads containing SAGs were then specifically isolated into multi-well plates for re-amplifying as storage SAG libraries (Supplementary Fig. 1a). The use of enzyme cocktails enhances the genome cover rate by 9.1-25% and improves genome amplification biases compared to conventional alkaline lysis treatment (Supplementary Fig. 1b and c). We confirmed that SAG-gel excludes the risk of cross-contamination between gel beads, showing all of the sequenced SAGs had > 99.6% of their reads mapping to either *E. coli* or *B. subtilis* (Supplementary Fig. 1d). The de novo assembled contigs showed a lower number of misassembled and unaligned contigs in SAG-gel (Supplementary Fig. 1e and Supplementary Table 1). In addition, SAG-gel showed a larger length of N50 and lower number of mismatches and indels, suggesting that accurate genome information can be acquired compared to conventional in-tube WGA reactions.

### Single-cell genome sequencing from a variety of environmental bacteria

We applied SAG-gel for single-cell genome sequencing of 8 environmental samples including 6 soil samples (beach soil: S1; desert soil: S2; mangrove soil: S3; fresh sea sediment: S4; frozen sea sediment: S5; and seashore soil: S6) and 2 seawater samples (harbor seawater: W1 and open ocean seawater: W2). At first, we individually generated 1,970 SAG reactions from isolated gel beads and 1,031 out of them were identified as positive prokaryotic fractions according to PCR and 16S rRNA gene sequencing. Marine microbiome (S5 and S6) showed higher positive reaction rates (> 61%) than soil microbiome (29-56%). After the sequencing of 929 positive prokaryotic fractions and subsequent data curation, we recovered 16 high-quality, 244 medium-quality, and 474 low-quality draft genomes, ranging from 0.65 to 5.8 Mbp in total length and 18 to 1,347 contigs (Fig. 2 a, b, Supplementary Table 2). High-quality SAGs yielded an average N50 of 83.2 kb, while medium- and low-quality ones yielded 27.5 kb and 12.3 kb, respectively. Regarding the presence of tRNAs from high-to-low SAG quality, the average numbers were 19.8, 16.5, and 10.6, respectively. The SAG-containing beads isolation with FACS effectively prevented the contaminated SAG prevalence (7.62% (56/735)) compared to the bead isolation with manual picking (38.7% (75/194)) (Supplementary Fig. 2). The average completeness for each sampling site was 32.9% in S1, 37.1% in S2, 42.3% in S3, 36.2% in S4, 54.8% in S5, 39.4% in S6, 42.2% in W1, and 53.6% in W2 (Fig. 2 c), while the completeness was broadly distributed from 1.37% to 99.9%.

**Figure 2.**
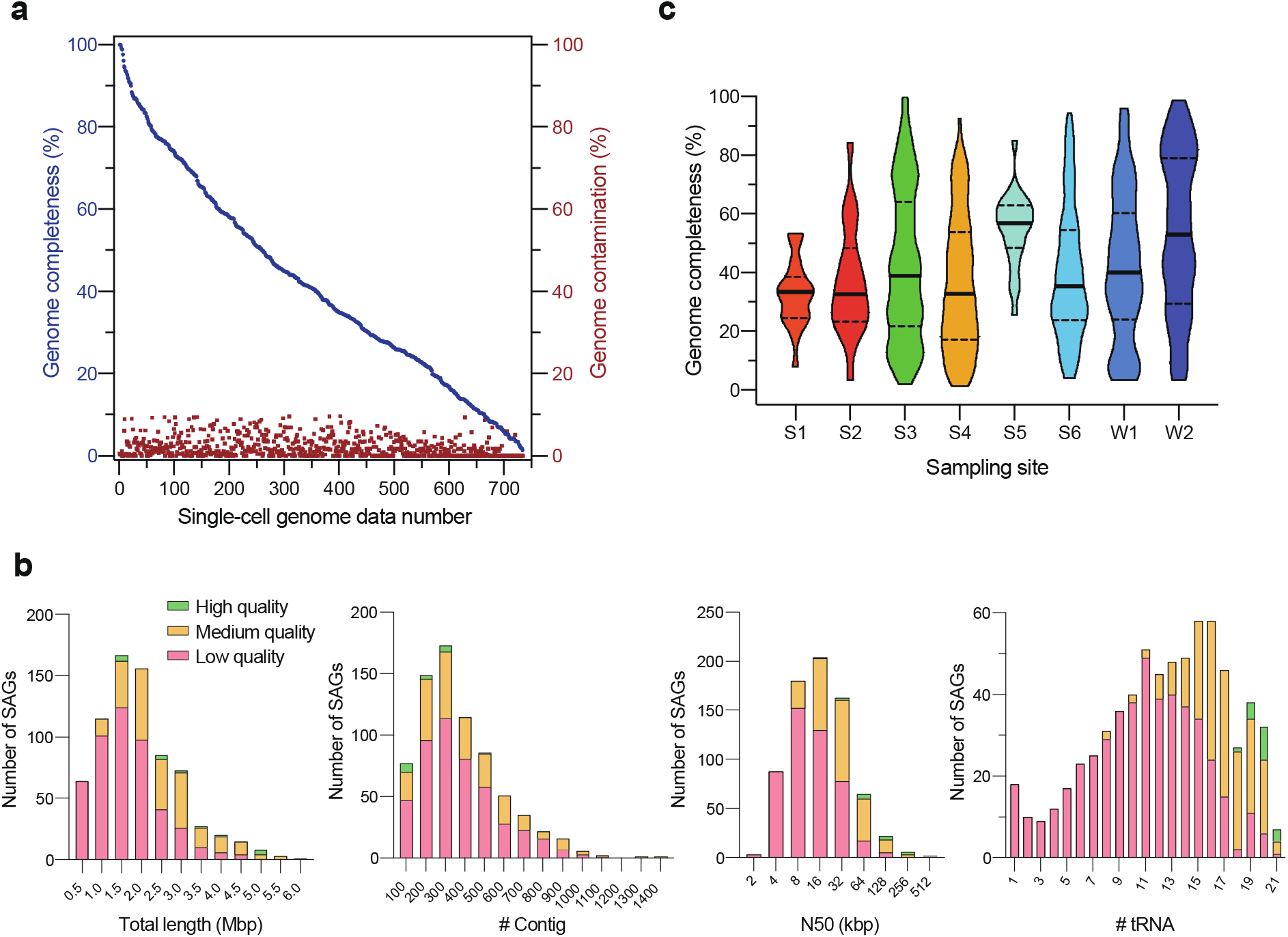
734 SAGs recovered from 8 environmental samples. a: Genome completeness and contamination of each SAG is plotted. The SAGs were classified as high, medium, low, and contamination according to the score of completeness and contamination, and the presence of tRNA sequence detected defined by the Genomic Standards Consortium. b: Distributions of the number of contigs, the number of tRNA, N50, and total length classified by the quality of draft genomes. c: Distributions of completeness in each sampling site. Solid line is the median and dotted line is the quartile.

### Taxonomic distribution of SAGs

Out of 734 SAGs, 76.0% (558/734) recovered > 500 bp 16S rRNA gene sequence and 58.7% (431/734) were > 1,400 bp in length (Supplementary Table 3). SAG-gel platform yielded longer sequences for genome assemblies and notably higher recovery rates of 16S rRNA gene compared to 7–17% in metagenome-based analysis^5, 14^ and 23–27% in other single-cell genome analyses^15, 16^. The 16S rRNA gene-based phylogenetic analysis revealed that 117 (102 specific) sequences > 1,400 bp and 204 (173 specific) sequences > 500 bp have no obvious reference sequences (Fig. 3a). To annotate all SAGs including those without 16S rRNA gene sequence, we utilized GTDB-Tk^17^ for taxonomic classification, resulting in 650 SAGs classified as 11 archaea and 639 bacteria, consisting of 44 phyla, 75 classes, 117 orders, and 127 families (Supplementary Table 3). *Proteobacteria* was dominant [277 (43.3%)] and *Desulfobacterota* [70 (10.9%)] and *Cyanobacteriota* [44 (6.9%)] followed. We found a total of 231 species from 650 SAGs (Supplementary Table 4), 98.7% (228 species) of which were newly identified with > 0.05 mash distance to reference genomes registered in Refseq^18^, suggesting that SAG-gel enables the large-scale accumulation of undescribed genomes from complex microbial communities (Supplementary Fig. 3).

**Figure 3.**
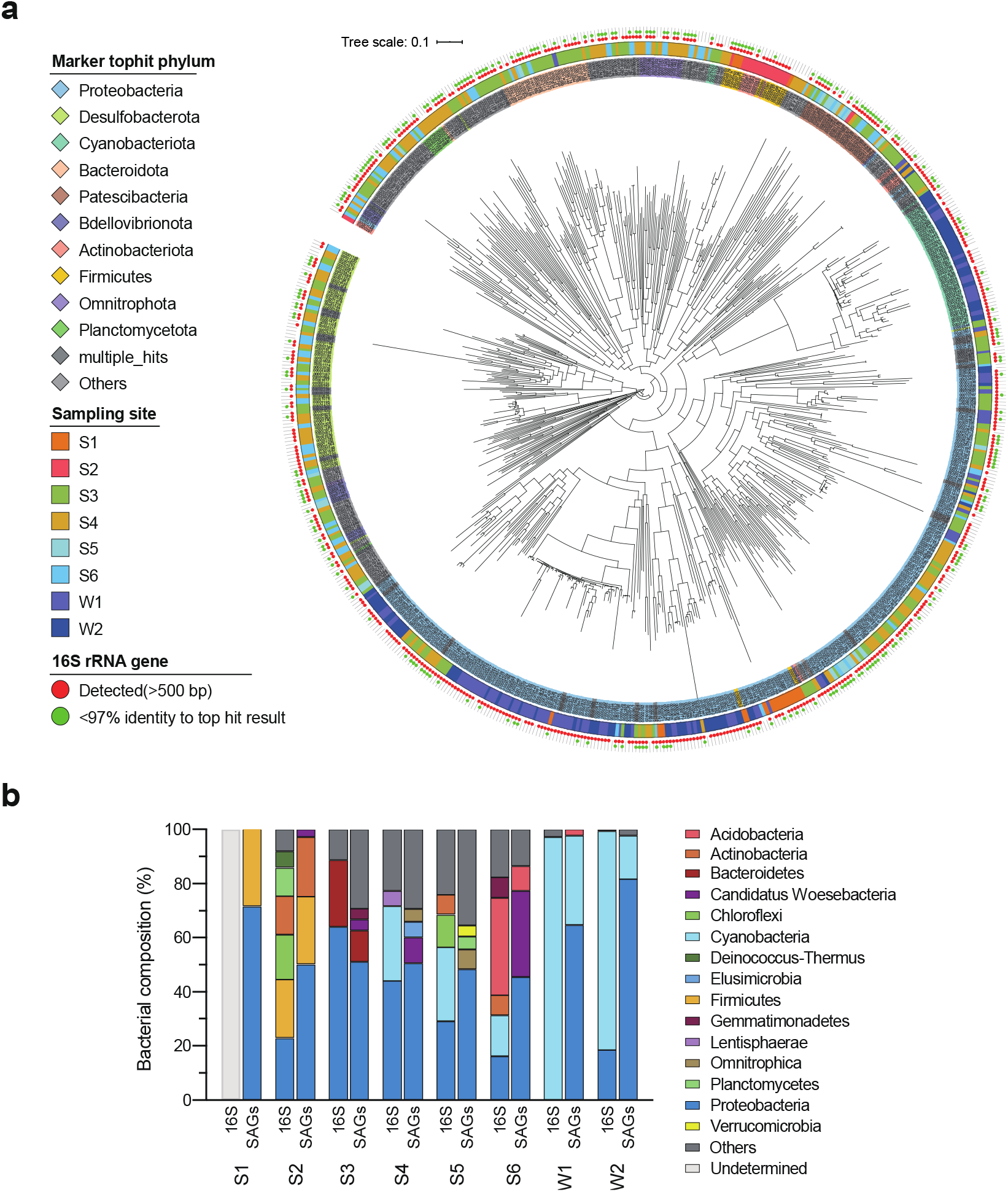
Taxonomical classifications of SAGs. a: Taxonomic annotation of 639 bacterial SAGs by phylum level. Inner circle: Top hit phylum; outer circle: sampling site; inner dot: 16S rRNA gene detected; outer dot: < 97% identity with top hit result. Any phyla which was not in the top ten phyla was clustered as “others.” b: Comparison of bacterial composition in 16S rRNA gene sequencing and SAGs (Phylum level). Taxonomic identification of SAGs was conducted by GTDB-Tk. In S1, the bacterial composition of 16S rRNA gene sequencing was undetermined because of the shortage of extract DNA. Any phyla which shared < 5% was clustered as “others”.

We compared the phylum-level taxonomic compositions assessed from SAGs and metagenomic 16S rRNA gene amplicon sequencing (Fig. 3b). While the number of major phyla (> 5% in relative abundance) detected in 16S rRNA gene sequencing was 11, SAGs recovered 10 phyla—all except for *Chloroflexi*, which is reported to be a tough bacterium for DNA extraction^19^. The bacterial composition of SAGs was not consistent with the composition of 16S rRNA gene sequencing except for mangrove (S3), where the result also corresponded to the previously reported data^20^ (Supplementary Fig. 4). In general, because SAGs reflect the exact cell numbers whereas 16S rRNA gene sequencing reflects the copy number of 16S rRNA genes and is also affected by amplification bias in the library preparation step^21^, these two methods may exhibit different bacterial composition. From the randomized sampling, SAG-gel can also recover SAGs of very rare phyla (< 0.1% in 16S rRNA gene sequencing), including *Omnitrophota* bacteria in sea sediment (S4) and *Elusimicrobiota* bacteria in seashore (S6).

### Determining gene distributions at single-cell resolution

From each sample set consists of 734 SAGs, we compared sampling site-specific enriched genes. Out of 3,221 orthologous gene groups (OGs)^22^ consisting of > 100 genes, 2,621 OGs (81%) were enriched (p < 0.01) in at least one sampling site (Fig. 4a) (Supplementary Table 5). For example, OGs related to phosphate acquisition (OG0000970, 0002021) were enriched in seawaters (W1 and W2) as previously reported^23^. In addition, phycobilisome protein (OG0001535) and phycobilisome linker polypeptide (OG0001628) were enriched in W2, while cytochrome c (OG0000546) and cytochrome P450 (OG0000546) were enriched in W1. In mangrove (S3) where heavy metals are accumulated^24^, OGs related to heavy metal resistance (OG0001092) exhibited higher abundance. In sea sediment (S4), sulfatase (OG0000005) and the radical SAM superfamily (OG0000056), which are mainly derived from *Omnitrophota* and *Desulfobacterota* bacteria, were enriched, while these features were not detected in frozen sea sediment (S5). In desert (S2), OGs related to spore germination (OG0002927 and OG0003158) were enriched. In seashore (S6), OGs related to aldehyde ferredoxin oxidoreductase (OG0000150) mainly derived from archaea were enriched. These results reflect the characteristics of each sampling site or sample storage conditions. The extraction of genomic features from a set of multiple draft genomes is useful for overviewing the whole microbial communities of the target environment like a metagenomic analysis manner. Furthermore, we can reveal which bacteria are responsible for the environment-specific genetic characteristics at single-cell resolution by in-depth sequence assignments.

**Figure 4.**
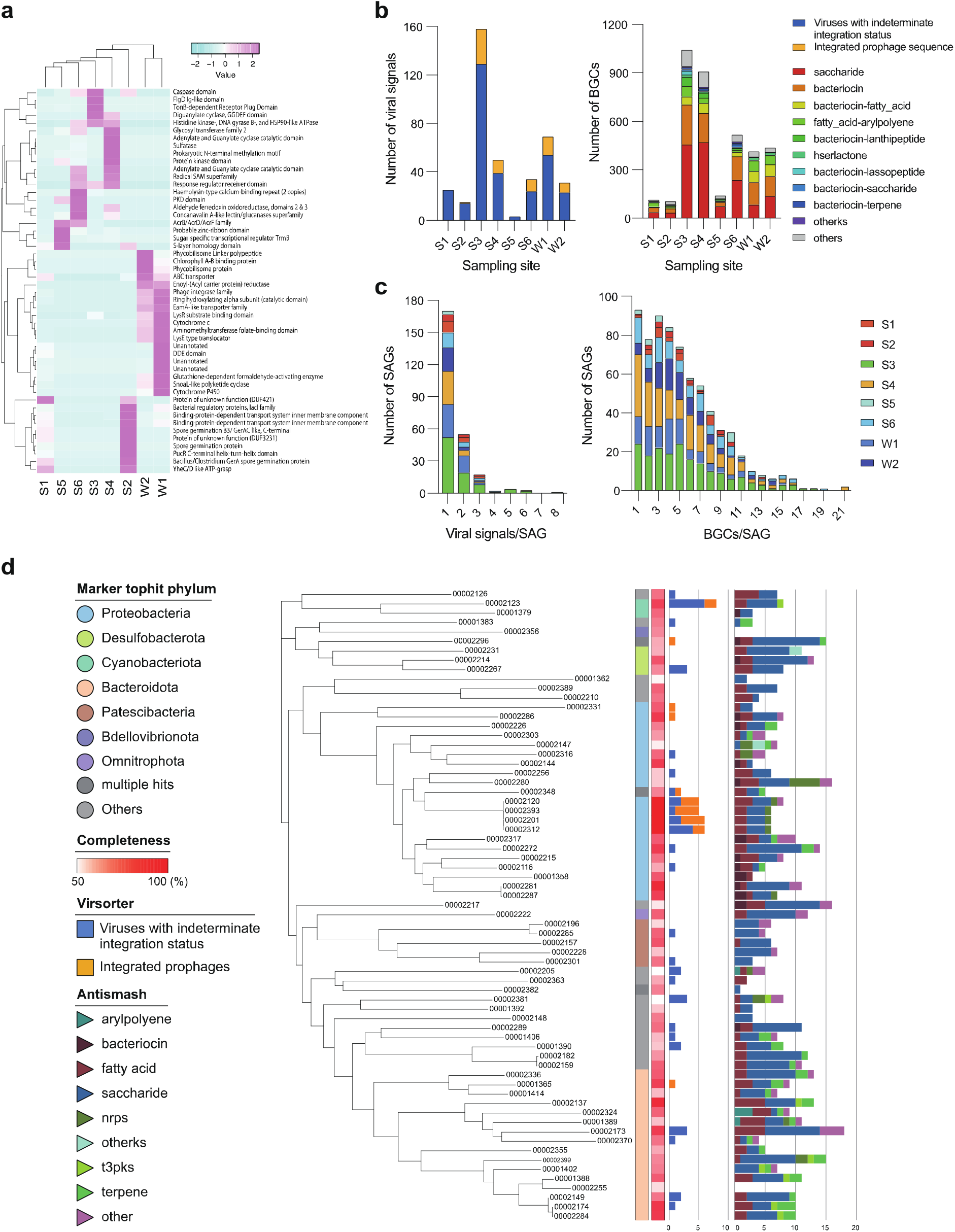
SAG-gel reveals distributions of viral signals and secondary metabolite biosynthetic gene clusters (BGCs) in target environment. a: Heatmap of sampling site specific orthologous gene groups (OGs) (p < 0.01) based on the Z score. Out of 3221 orthologous gene groups (OGs) which consist of > 100 genes, 2621 OGs were enriched (p < 0.01) in at least one sampling site and the smallest 50 OGs were extracted for figure drawing. b: Detected gene number of viral signals and BGCs classified by their category. In VirSorter, phages matching the prophage database (VirSorter categories 1 and 2: viruses with ‘indeterminate integration status’; 4 and 5: integrated prophage) were included. In antiSMASH, BGCs of detected numbers <10 were classified as “others.” c: Distributions of viral signals and BGCs by SAGs colored by sampling sites. d: Completeness, taxonomical classification, and the number of viral signals and BGCs were summarized by SAGs in S3: mangrove soil. High- and mediumquality SAGs were included. In the result of BGCs, clusters assigned as “cf_putative” were not counted.

Viral signals and biosynthetic gene clusters (BGCs) were detected from various kinds of bacteria in each sampling site, suggesting that these genes are widely spread in the area surrounding the Red Sea (Fig. 4b, c) (Supplementary Table 6, 7). In contrast, most plasmid sequences were distributed in *Enterobacteriaceae* (Supplementary Table 8), which may be due to the bias of the existing database^6^. In terms of viral sequences, 385 sequences were detected in total, with about 34.3% (252/734) of SAGs carrying at least one viral sequence. In soil samples, 77% (17/22) of SAGs carried at least one viral sequence in beach (S1), followed by desert (S2) (53%) and mangrove (S3) (44%). In contrast, seawaters (W1 and W2) exhibited lower rates (20 and 13%, respectively). When we focused on viral signals per single cell, a SAG from mangrove (S3) assigned as *Cyanobacteria* bacterium UBA9579 carried 8 viral sequences while the median number of viral signals in single cells was 1.0 (Fig. 4c). On the other hand, 3,676 BGCs were detected in total, with about 94.6% (694/734) of SAGs carrying at least one BGC. Saccharide was dominant (41.5%), followed by bacteriocin-related BGCs and fatty acid-related BGCs (Fig. 4b). While the median number of BGCs in single cells was 5.0, two SAGs assigned as *Acidobacteriota* bacterium UBA890 and *Oligoflexales* bacterium from sea sediment (S4) had 21 BGCs (Fig. 4c). In *Acidobacteriota* bacterium UBA890, 8 bacteriocins were detected. In *Oligoflexales* bacterium, 7 saccharides were detected (Supplementary Fig. 5). SAG-gel links the taxonomy and metabolic functions at single-cell resolution, thus enabling to search for the target genes or target hosts from complex microbiome on demand, which is difficult in metagenomic binning approaches (Fig. 4d).

### Whole-genome comparative analysis of bacteria within species identical in 16S rRNA gene

From seawater (W1), we found 28 SAGs shared 99.9% identities over 1,457 bp of 16S rRNA gene and assessed the average nucleotide identity (ANI) and ortholog presence/absence within them. They were assigned as *Rhodobacter* spp., which are a commonly found genus in freshwater or marine environments and are known to have a wide range of metabolic capabilities^25^. ANI suggested that 28 SAGs fall into several clusters, with 2 large clusters (*Rhodobacter* spp. RS1 and RS2) (Supplementary Fig. 6a). The intra-cluster ANI was 95.3 (RS1) and 96.3 (RS2) while the inter-cluster ANI was 91.1. We then combined each SAG^26^ and two draft genomes of RS1 and RS2 were acquired that had not been described in the previously reported catalogue of microbial draft genomes from the Red Sea^27^. The completeness and contamination of RS1 and RS2 were 80.16% and 2.19%, and 91.39% and 4.33%, respectively. Both are > 2.8 Mb in genome size with > 2,900 coding DNA sequences (CDSs). The sequence identity of 5S rRNA (109 bp) and 23S rRNA (2,558 bp) was 100% and 99.8%, respectively. While 143 draft genomes of 20 species including uncultured *Rhodobacter* have been registered in the NCBI taxonomy database, RS1 and RS2 most closely related to *Rhodobacteraceae* bacterium HIMB11 isolated from coastal seawater in Kaneohe Bay^28^, exhibiting > 99.6% identity with 5S, 16S, and 23S rRNA sequences. Although the genome sizes and CDS (3.1 Mbp and 3,183 in HIMB11) are comparable to each other, ANI over the whole genome was 91.7 (RS1-RS2), 91.4 (RS1-HIMB11), and 92.4 (RS2-HIMB11), respectively. When we compared the average amino acid identity (AAI) of the translated CDSs, the score was 91.4% (RS1-RS2).

*Rhodobacter* spp. RS1 and RS2 diverged genetically and showed different alignments in the entire genomic sequences while retaining the high sequence identity of the 16S rRNA gene (Supplementary Fig. 6a, b). We compared the sequences of 13 draft genomes including 6 from RS1, 6 from RS2, and HIMB11, which suggested that each strain shared the unique gene sets (Fig. 5). In total, 34,423 genes classified into 6,274 clusters were defined and the genes belonging to only one SAG constituted 6.9% (2,372/34,423), 92.8% (2,202/2,372) of which were unannotated. KEGG pathway analysis^29^ with HIMB11 revealed that the module completion ratio (MCR) of glucose/mannose transport system and fructose transport system was 25% in RS1 and RS2, while the MCR of phosphonate transport system, iron complex transport system, zinc transport system, putative zinc/manganese transport system, and lipopolysaccharide export system was 100% only in RS1 and RS2 (Supplementary Table 9). SAG-gel clarified the incongruence of the 16S rRNA gene and ANI within some *Rhodobacter* spp. and identified the different functional modules. This single-cell resolution genome analysis can shed light on these taxonomic and metabolic incongruences obscured in uncultured bacteria and provide highly-resolved microbial diversities^30^.

**Figure 5.**
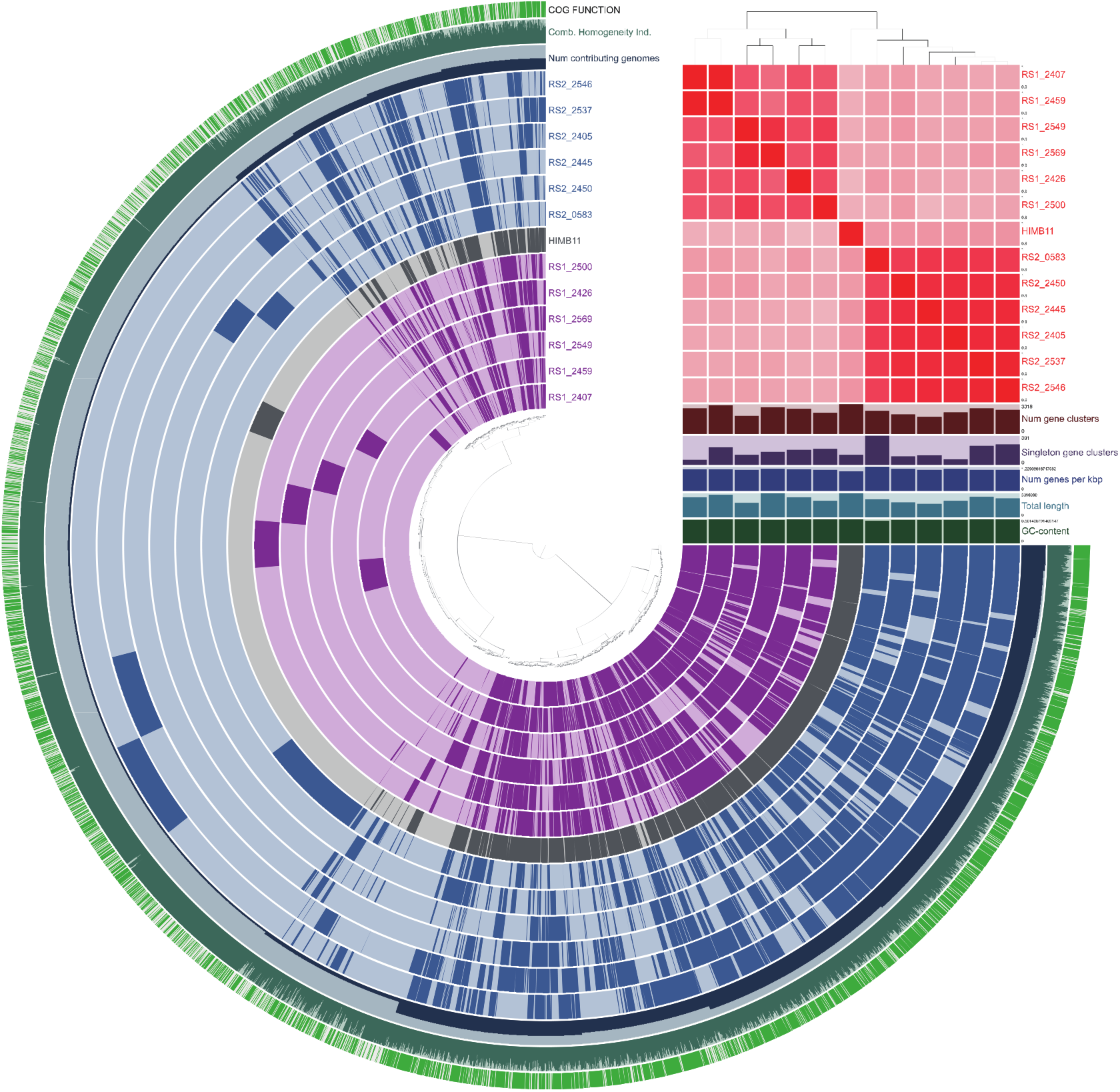
Comparative genome analysis of *Rhodobacter* spp. Comparative genome analysis of 6 RS1 SAGs and 6 RS2 SAGs. HIMB11, the reference genome, is the genome of known *Rhodobacter* sp. containing > 99% identical 16S rRNA gene. The figure shows the presence-absence of 6,274 gene clusters in the pangenome of 12 SAGs and the reference genome. Inter-strain average nucleotide identity showed < 94%, and RS1-specific or RS2-specific gene clusters were detected.

## Discussion

Unlike metagenomic binning or the conventional single-cell genomics, SAG-gel associated single-cell genomics efficiently provides draft genomes containing fulllength 16S rRNA genes from the complex microbial communities. In SAG-gel platform, the SAG-containing beads can be stored for months at 4°C and the plate-based SAG library can be frozen for longer storage, expanding the possibility to conduct scaled-up sequencing in various sequencing machine on demand. Although the randomized single-cell collection manner is feasible to accumulate the genetic information according to the real microbial abundance in the environment, we would be able to enrich specific microbial fraction with the sample preprocessing such as cell size fraction and the specific cell enrichment based on phylogenetic or functional defining combining with fluorescent markers with FACS. In addition, prior to deep sequencing, the SAG library can be screened for specific genes or taxa based on marker genes at single-cell resolution, which will be useful for identifying pathogenic bacteria and metabolic producers by detecting viral signals, BGCs, and plasmids even when the target cell is underrepresented.

We demonstrated SAG-gel can be applied to diverse environmental samples including soil and seawater without any modification of protocols because of its versatility. From the first screening of prokaryotic amplicons from SAG reactions, about half of SAGs were successfully obtained while showing the presence of the 16S rRNA gene. The yield ratio for uncontaminated SAGs was 79% (734/929) which includes rare bacteria in 16S rRNA gene sequencing. By using FACS for fluorescence-positive beads isolation, 84% of SAGs were assigned as draft genomes with > 0% completeness and < 10% contamination. FACS-based beads isolation is feasible for the selection of SAGs from the pool of amplified DNA containing beads in high throughput and contamination-less manner (Supplementary Table 2). The combination of the randomized SAG-gel pool and FACS-based beads isolation enables us to obtain bias-free genomic population data from individual cells and provide an overview of the environment-specific genetic potential like metagenomic population analysis (Fig. 4). It also revealed that there are a variety of microbial species in the environment that have not been described yet (Fig. 3). The newly obtained SAGs were sufficiently detailed to be described and be compared their gene characteristics at the single-cell resolution (Supplementary Fig. 5).

The current metagenomic approach is powerful to estimate the environment-specific gene contents; furthermore, the metagenomic binning tools enable the production of draft genomes based on nucleotide composition and coverage depth. However, when multiple species or strains belonging to the same genera are present at similar abundances and have similar nucleotide compositions, the metagenomic binning approach often fails to separate contigs from these taxa into the correct genome bins^31^. In contrast, our approach generates individual SAGs for each species, which can be clustered by pairwise determination of ANI or taxonomic marker identities. As shown in the comparative genome analysis of uncultured *Rhodobacter* spp., massive SAG production reveals the intra-species diversity in individual environments. We found that the co-existence of multiple *Rhodobacter* strains in the same microbial community and they presumably have different functional modules while retaining highly conserved 16S rRNA gene sequences (Fig. 6 and Supplementary Fig. 6). We should pay close attention to the existence of these taxonomically identical but functionally different microbes, especially in the sequencing of geographically separated environmental samples and symbiotic or commensal bacteria from different hosts.

Our results reveal that the single-cell draft genomes work as *de novo*-assembled sequences as well as the genomes of isolates. However, unlike the genomes of isolates, challenges in genome assembly are not fully resolved using SAG-gel platform alone, as WGA process generates numerous chimeric reads^32^, hampering the assembly of long contiguous sequences and randomly biased amplified genomes^11^, and causing incomplete genome coverage. By combining our methods with other tools including cleaning and co-assembly of SAGs from the same species, the number of chimeric contigs and genome incompleteness could be significantly reduced^26, 33^. Draft genomes acquired by SAG-gel can also be combined with other sequence data sets including shotgun metagenome sequencing and long-read sequencing, all of which will promote genome finishing^34^. Moreover, we recognize the shortage of sequenced SAGs to cover most microbial species in complex microbial communities. In our estimation, we could pool samples as 384- to 768-plex SAG libraries for sequencing in a single lane on the Illumina HiSeq platform while acquiring enough sequencing depth for *de novo* assembly.

The ability of SAG-gel to generate qualified draft genomes in a high-throughput manner will contribute to increases the comprehensiveness of reference genomes without the need for laborious isolation and cultivation. While we focused on the environmental microbiome, SAG-gel platform is also applicable to other types of cells including eukaryotic cells. We expect that SAG-gel platform will contribute to disclose the diversity and heterogeneity of biological communities.

## Methods

### Fabrication of microfluidic device

A microfluidic device for generating monodispersed picoliter-sized droplets was fabricated according to a previously-reported method^35^. In short, a flow-focusing microfluidic device was designed using AutoCAD (AutoDesk, Sausalito, CA, USA) and fabricated using conventional soft-lithography techniques. A photomask pattern was transferred to a layer of negative photoresist (SU8-3050; Microchem, Newton, MA, USA) coating a glass wafer (40 mm × 47 mm). All microchannels were 50 μm tall and 100 μm wide, except at the cross-junction area. The cross-junction was designed to be 17 μm wide for the aqueous phase and 10 μm wide for the continuous oil phase. Polydimethylsiloxane (PDMS; Sylgard 184; Dow Corning Corp., Midland, MI) and its cross-linker were mixed thoroughly at a ratio of 10:1 (w/w) then degassed. The PDMS mixture was poured over the master mold and cured for at least 1 h at 60 °C. After curing, the slabs were punched with a 0.75-mm biopsy punch (World Precision Instruments, Sarasota, FL, USA) and PDMS slabs and glass slides were bonded by plasma treatment (Plasma Cleaner PDG-32G; Harrick Scientific, Ossining, NY, USA) followed by baking for at least 30 min at 60 °C. Finally, the microchannel was filled with Aquapel solution (PPG Industries, Pittsburgh, PA, USA) to produce a hydrophobic surface coating; thereafter, excess Aquapel was blown off with air.

### Preparation of model bacteria suspensions

For genome sequencing analysis, *E. coli* K-12 strain (ATCC 10798; genome size: 4.6 Mbp) and *B. subtilis* (ATCC 6633; genome size: 4.0 Mbp) were used as model bacteria. *E. coli* K-12 cells were pre-cultured in Luria-Bertani (LB) medium (1.0% Bacto tryptone [BD Biosciences, Franklin Lakes, NJ, USA], 0.5% yeast extract [BD Biosciences], 1.0% NaCl [Sigma-Aldrich, Hamburg, Germany], pH 7.0) for 16 h. *B. subtilis* cells were pre-cultured in brain heart infusion broth (ATCC medium 44; Thermo Fisher Scientific, Waltham, MA, USA.) for 16 h. For cell collection, 1 mL of cultured medium was dispensed into a 1.5 mL tube and centrifuged at 8000 × *g* for 5 min. After removing the supernatant, the collected cells were resuspended in UV-treated Dulbecco’s phosphate-buffered saline (-) (DPBS, Thermo Fisher Scientific) and washed three times with DPBS. Finally, the cells were resuspended in 500 μL DPBS and cell concentration was calculated with a bacteria counter. To evaluate cross-contamination, *E. coli* and *B. subtilis* cells were mixed at a ratio of 1:1. For single-cell encapsulation, the cell concentrations of *E. coli* and *B. subtilis* were adjusted to 3.0 × 10^3^ cells/μL at a concentration of 0.1 cell/droplet. All preparations for the cell suspension and further processes were performed under an open-interior clean bench (KOACH T 500-F; KOKEN LTD., Tokyo, Japan) except for droplet generation and isolation with FACS.

### Environmental sample preparation

Eight samples from seven different sampling sites were collected for this study. Environmental samples consisted of six soil and sediment samples [beach soil: S1(22°17’35.6”N 39°05’26.0”E); desert soil: S2(22°19’03.0”N 39°08’36.7”E); mangrove soil: S3(22°18’53.3”N 39°05’29.2”E); fresh sea sediment: S4(22° 17.988’N, 39° 03.427 ’E); frozen sea sediment: S5(22° 17.988’N, 39° 03.427 ’E); and seashore soil: S6(22°17’17.5”N 39°05’42.3”E)] and two seawater samples [harbor seawater: W1(22°18’16.9”N 39°06’12.3”E) and open ocean seawater: W2(22° 17.988’N, 39° 03.427 ’E)]. For soil samples, 20–30 g of soil was collected 10 cm beneath the top layer and maintained in 50 mL tubes on ice until preparation of the cell suspension. For sediment samples, 20–30 g of soil was collected from the seafloor at the depth of 25 m with van Veen Grab sampler and kept into 50 mL tubes. For the S5 sample, sea sediment was collected on 30th April 2017, frozen by dry ice, and directly kept on −80°C for 14 months. It was thawed on ice immediately before the preparation of cell suspensions. S4 was also kept on ice until the preparation of cell suspensions. W1 and W2 was collected from the surface. From each sampling site, 30 L of seawater was collected into plastic tanks. Cell suspensions were prepared as soon as possible after arrival at the laboratory. All samples except for S5 were collected from the 24th to the 27th of June, 2018 and proceeded to WGA.

Ten grams each of samples S1 to S6 was dispensed into three 50 mL tubes (Iwaki Science Products Department, Iwaki Glass Co. Ltd., Chiba, Japan). Then DPBS was added to each tube up to the 40-mL volume marker and the contents were mixed thoroughly. The suspended solution was kept on ice for 5 min and the supernatant was collected into another 50 mL tube. The supernatant was filtered with a 5-ψm MF-Millipore membrane filter (Merck Millipore, Milan, Italy). The flow-through was centrifuged at 10,000 ×*g* for 5 min with a benchtop centrifuge (Thermo Fisher Scientific). After removing the supernatant, the pellet was resuspended in 10 mL of DPBS and dispensed into a 1.5 mL tube (Axygen Biosciences, Hangzhou, China). Each suspension was centrifuged at 10,000 ×*g* for 5 min with a tabletop ultracentrifuge and washed three times with DPBS.

For the seawater samples, 4 L of seawater was filtered with a 5-ψm MF-Millipore membrane filter. The flow-through was collected and filtered with a 0.22-μm filter (Merck Millipore). Then the 0.22-μm filter was suspended in 10 mL of DPBS and vortexed thoroughly to suspend trapped bacterial fractions; 10 mL of bacterial suspensions was dispensed into 1.5 mL tubes, centrifuged at 10,000 ×*g* for 5 min with a tabletop ultracentrifuge, and washed three times with DPBS.

We prepared two tubes of cell suspensions for each sample. One tube was used for single-cell genome sequencing and the other for 16S rRNA gene sequencing. For single-cell genome sequencing, the cell concentration was adjusted to 3.0 × 10^3^ cells/μL for encapsulating single cells at a concentration of 0.1 cell/droplet (40 μm diameter). For 16S rRNA gene sequencing, total metagenomic DNA was extracted with PowerLyzer Soil DNA Extraction kit (QIAGEN) from the cell suspensions. Then 16S rRNA gene sequencing libraries were prepared according to Illumina’s protocol and run on an Illumina MiSeq for 300 cycles of paired-end sequencing using a MiSeq v3 600-cycle reagent kit (Illumina, San Diego, CA, USA). For S1, the amount of DNA extracted was insufficient to prepare the sequencing library.

### Single-cell encapsulation into agarose gel beads

Ultra-low gelling temperature agarose A5030 (Sigma-Aldrich) was mixed into DPBS and incubated at 85 °C for 30 min. By mixing the cell suspensions with agarose solutions, 1.5% agarose cell suspensions with 3.0 × 10^3^ cells/μL were prepared. Agarose cell suspensions were loaded into PTFE tubing (AWG 24) connected to a Mitos P-pump (Dolomite, Charleston, MA, USA). By controlling the air pressure, agarose cell suspensions and 2% Pico-Surf™ 1 in Novec™ 7500 (Dolomite) as carrier oil were introduced into a manufacturer-fabricated microfluidic device. Microfluidic droplets with a diameter of 40 μm (volume: 34 pL) were generated for encapsulation of > 100,000 single cells within 30 minutes (35,000 droplets/min) at a concentration of 0.1 cell/droplet.

Droplets were collected in 1.5 mL tubes via PTFE tubing from the outlet and incubated on ice for 15 min. After droplet solidification, the collected droplets were broken by 1H,1H,2H,2H-perfluoro-1-octanol (Sigma-Aldrich). Next, 500 μL of acetone (Sigma-Aldrich) was added to the 1.5 mL tube and vortexed thoroughly. Gel beads descended to the bottom of the 1.5 mL tube because of their greater specific gravity, which enables collection of gel beads by centrifugation. After collecting the gel beads with a tabletop centrifuge, the supernatant was removed. Then 500 μL of isopropanol (Sigma-Aldrich) was added to the 1.5 mL tube and vortexed thoroughly. After gel-bead collection with a tabletop centrifuge, the supernatant was removed. Finally, 500 μL of DPBS was added to the 1.5 mL tube and vortexed thoroughly. The gel beads were washed with DPBS three times. During these steps, the gel beads were transitioned from an oil phase to an aqueous phase.

### Cell lysis and WGA in agarose gel beads

After the gel beads were collected with a tabletop centrifuge and supernatant was removed, they were treated with two different lysis protocols: Alkaline treatment and Enzyme cocktail treatment.

#### 1) Alkaline treatment and WGA

The following steps were conducted in 0.2 mL tubes (Axygen Biosciences): 3 μL of D2 buffer from a REPLI-g Single Cell Kit (Thermo Fisher Scientific) was added to 4 μL of droplet suspension. After incubation at 40 °C for 10 min, 40 μL of WGAm mixture (3 μL of Stop Solution, 9 μL of H_2_O, 29 μL of reaction buffer, and 2 μL of DNA polymerase) was added and incubated at 30 °C for 180 min.

#### 2) Enzyme cocktail treatment and WGA

Two hundred microliters of lysozyme solution (50 U/μL Ready-lyse lysozyme [Lucigen, WI, USA], 2 U/mL zymolyase [Zymo Research, Orange, CA, USA], 22 U/mL lysostaphin [Sigma-Aldrich], and 250 U/mL mutanolysin [Sigma-Aldrich] in DPBS) was added and incubated at 37 °C overnight. Gel beads were washed with DPBS three times and 200 μL of achromopeptidase solution (0.5 mg/mL achromopeptidase [Wako, Tokyo, Japan] in DPBS) was added and incubated at 37 °C for 8 h. Then the gel beads were washed with DPBS three times and 200 μL of proteinase K solution (1 mg/mL proteinase K [Promega, Madison, WI] and 0.5% SDS [Wako, Tokyo, Japan] in DPBS) was added and incubated at 40 °C overnight. After washing the gel beads with DPBS five times, the supernatant was removed. Then the gel beads proceeded to the alkaline treatment and WGA described above.

For SAG-gel with model bacteria, both two lysis protocols were performed for evaluation, while enzyme cocktail treatment was performed for environmental samples.

### Confirmation of DNA amplification

After WGA, the gel beads were collected with a tabletop centrifuge and the supernatant was removed. The gel beads were resuspended in DPBS and washed three times. Then the gel beads were stained with 1× SYBR Green (Thermo Fisher Scientific) in DPBS and transferred onto a glass slide for microscopic observation. After confirmation of DNA amplification, fluorescence-positive gel beads were isolated by FACS or manual picking. For the model bacteria experiment, gel-bead isolation was conducted by FACS. For the environmental sample experiment, gel beads from S1, S2, S5, S6, and W1 were subjected to manual picking within 2 days after WGA. The remaining samples were maintained in DPBS and isolated by FACS within 1 week. To compare the isolation methods, both manual picking and FACS-based isolation were conducted for S6 and W1. From 8 sampling sites, 48–480 fluorescence-positive gel beads were isolated and subjected to 2nd-round WGA.

#### 1) Single-droplet sorting with manual picking

SYBR-stained gel beads were transferred onto a glass slide. Fluorescence-positive gel beads were manually picked using a micropipette (Drummond, Camlab, Cambridge, UK) under the open clean system (KOKEN LTD.). Then each bead was dispensed into a 96-well plate with 0.8 μL of DPBS and proceeded to a 2nd-round of WGA or maintained at −30 °C for longer storage.

#### 2) Single-droplet sorting with FACS

Flow cytometric analysis and sorting were performed with a BD FACSMelody™ Cell Sorter (BD Biosciences) equipped with a 488 nm excitation laser. The nozzle diameter was 100 μm. The sample flow rate was adjusted to approximately 30 events per second. The number of sorted beads is summarized in Supplementary Table 1a, b; 0.8 μL of DPBS was dispensed into each well prior to gel-bead sorting. After sorting, the 96-well plate proceeded to a 2nd-round of WGA or maintained at −30 °C for longer storage.

### 2nd-round WGA

2nd-round WGA was performed with the REPLI-g Single Cell Kit (Thermo Fisher Scientific); 0.6 μL of D2 buffer was added to each well and incubated at 65 °C for 10 min. Then 8.6 μL of WGA mixture (0.6 μL of Stop Solution, 1.8 μL of H_2_O, 5.8 μL of reaction buffer, and 0.4 μL of DNA polymerase) was added and incubated at 30 °C for 120 min. The WGA reaction was terminated by heating at 65 °C for 3 min. After the 2nd-round of amplification, master library plates of single amplified genomes (SAGs) were prepared.

### DNA quantification and 16S rRNA gene sequence analysis

The amplicon yields of the 2nd-round WGA were quantified by a Qubit dsDNA HS assay kit (Thermo Fisher Scientific). For model bacteria, 87 samples out of 92 (94.5%) showed sufficient DNA amplification for library preparation. For environmental samples, in order to confirm amplification from a single bacterial cell, 16S rRNA gene sequencing was also performed against the 2nd-round WGA products. Primer pair sequences for the V3V4 region were used according to Illumina’s MiSeq system protocols (Forward: 5’-TCGTCGGCAGCGTCAGATGTGTATAAGAGACAGCCTACGGGNGGCWG CAG-3’, Reverse: 5’-GTCTCGTGGGCTCGGAGATGTGTATAAGAGACAGGACTACHVGGGTA TCTAATCC-3’). PCR amplification was confirmed with agarose electrophoresis (100 V, 15 min) and the amplicon sequences were obtained via Sanger sequencing (Fasmac, Kanagawa, JAPAN).

### Construction of single-cell genome libraries and whole genome sequencing

For the sequencing analysis, Illumina libraries were prepared using amplicons from the 2nd-round WGA products. We used a Nextera XT DNA sample prep kit (Illumina) according to the manufacturer’s instructions. Libraries from model bacterial samples were sequenced on an Illumina MiSeq for 75 cycles of paired-end sequencing, generating a total of 50.0 M paired-end reads and 7.34 Gbp. Libraries from environmental samples were sequenced on an Illumina HiSeq for 150 cycles of paired-end sequencing (GENEWIZ, South Plainfield, NJ, USA), generating a total of 1.19 G paired-end reads and 356 Gbp.

### Sequencing analysis for model bacteria

The sequence raw reads were mapped to the NCBI reference genome (NIH, Bethesda, MD, USA) with BWA^36^ for evaluation of cross contamination. NC_00913 (*E. coli* substrain MG1655) and CP011496 (Escherichia coli strain NCM3722 plasmid F) were used as a reference for the *E. coli* K-12 strain. NCBI reference genome NC_014479 was used for *B. subtilis* subsp. spizizenii str. W23. Then the acquired reads were down-sampled to 10× mean mapping depth (46 Mbp for *E. coli* and 40 Mbp for *B. subtilis*). Sequence reads were *de novo* assembled with SPAdes 3.5.0^37^ and qualified by QUAST 2.3.^38^ Contigs (> 500 bp) from each sample were mapped to the reference genome with BWA. Genome coverage was calculated using SAMtools^39^.

### 16S rRNA gene amplicon sequencing analysis of environmental samples

Meta 16S short-read information of the V3V4 region was acquired from each environmental sample by Illumina MiSeq 300 cycle paired-end sequencing. OTU clustering was conducted by USEARCH^40^ from integrated pair-end reads longer than 400 bp and each OTU was annotated taxonomically by BLAST search against the Silva 132 database. OTUs were clustered by phylum level and the bar chart was drawn for the phyla accounting for > 5% (Fig. 3b).

### Sequencing analysis for environmental samples

Sequence reads were *de novo* assembled with SPAdes^37^ and contigs (> 1,000 bp) underwent the following analysis. Completeness and contamination were calculated with CheckM^41^. The number of contigs, N50, and total length was evaluated with QUAST^38^. The number of tRNA was examined with Prokka^42^. The draft genome quality was evaluated by the standards developed by GSC^43^. After removing samples which were classified as contamination or exhibited 0% completeness, the remaining SAGs underwent the following analysis. 16S rRNA gene sequence was extracted with Prokka and assigned to the 16S rRNA gene sequence database (Silva). The 16S rRNA genes which exhibited < 97% identity to the top hit result to the database were assigned to have no obvious reference sequences.

SAGs were taxonomically classified with GTDB-Tk^17^ and the results were visualized with iTOl^44^. Then the GTDB-Tk results were summarized at the phylum level to compare bacterial composition of SAGs and 16S rRNA gene sequencing. We evaluated the number of species using the mash distance 0.05 as the cutoff for species delineation, which equates to an ANI of ≥ 95 %^45^.

In order to characterize gene distributions at each sampling site, orthologous gene groups (OGs) were detected by OrthoFinder^22^. The detection number of each OG was summarized for each sampling site. Based on the number of OGs, the OGs with p-values < 0.01 were extracted. For the heatmap, 25 OGs exhibiting the lowest p-values were extracted.

SAGs were referenced against VirSorter^46^, antiSMASH^47^ (search option: -- transatpks_da --clusterblast --subclusterblast --knownclusterblast --smcogs --inclusive --borderpredict --full-hmmer) and Plasmidfinder^48^ for detecting viral signals, BGCs, and plasmids. For VirSorter, viral signals matching the VirSorter categories of 1, 2, 4, and 5 were considered and the viral signals matching the VirSorter categories of 3 and 6 were ignored. For antiSMASH, BGCs of detected numbers < 10 were classified as “others.”

SAGs that exhibited >95% ANI were co-assembled with ccSAG^26^ and the composite draft genomes and the draft genome of *Rhodobacteraceae* bacterium HIMB11 were evaluated with QUAST^38^, CheckM^41^, Prokka^42^, and FastANI^45^ (https://github.com/ParBLiSS/FastANI). Anvi’o^49^ was used to compare the sequences from 13 draft genomes (6 from RS1, 6 from RS2, and HIMB11) which were chosen in order based on the value of completeness. To visualize the alignments over the whole sequence, draft genomes of RS1 and RS2 were aligned to the shotgun sequencing data of HIMB11 with AliTV^50^. Filter links by their identity were 65–100% and filter links by their length were ≧ 2,500 bp. For detailed analysis, each draft genome was searched against Genomaple-2.3.2^29^ and module completion ratio (MCR) in each functional module was evaluated. Unless otherwise noted, the analysis was performed with default settings for each tool.

## Supporting information

Supplementary results

## Accession number

All sequencing data generated in this study will be available in the DDBJ Sequenced Read Archive under the accession numbers PSUB004727 upon final publication.

## Acknowledgements (optional)

This work was supported by JST-PRESTO grant number JPMJPR15FA, MEXT KAKENHI grant numbers 18H01801 and 17H06158, and the funding from King Abdullah University of Science and Technology (KAUST), under award numbers URF/1/1976/03/01, URF/1/1976/17/01, URF/1/1976/20/01 and FCS/1/3326/01/01. The super-computing resource was provided by the Human Genome Center (University of Tokyo).

## Author Contributions

YN, MK and MH contributed equally to this work.

YN, MK, MH, TG, and HT conceived and designed the experiments. YN, MK, MH, KT, and KM collected samples. HB conducted the DNA extraction. YN, MK, and KT conducted the other experiments and analyzed the results. YN, MH, MK, and HT wrote the manuscript.

