## Supplementary results for "Extensive single-cell genomics reveals bacterial diversity and diverse phage host ranges in the area in and around the Red Sea"

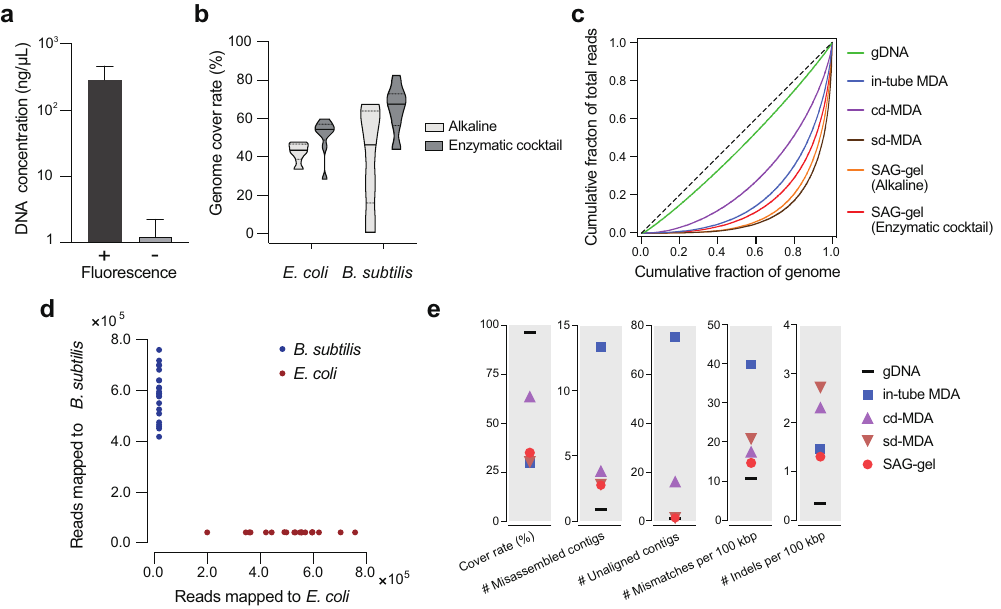

**Supplementary Figure 1. SAG-gel improves genome recovering of single bacterial cells.**

a: DNA concentration after 2nd-round WGA of fluorescence-positive and -negative beads

b: Comparison of genome cover rate of *E. coli* and *B. subtilis* treated with alkaline or enzymatic cocktail. Genome cover rate was evaluated at the ×10 sequence depth.

c: Lorenz curve for evaluation of amplification biases in single-cell sequence data

d: SAG-gel sequencing of mixtures of *E. coli* and *B. subtilis* cells. The scatter plot shows the number of reads mapped to *E. coli* and *B. subtilis* genomes associating to each droplet. Red dots indicate droplets that were identified as *E. coli* from these 16S rRNA data; Blue dots indicate droplets that were identified as *B. subtilis*.

e: Sequence statistics of contigs obtained from SAGs of *E. coli*.

In c and e, SAG-gel was compared with conventional in-tube MDA^1^ and previous reported method including compartmented droplet MDA (cd-MDA)^1^, single-droplet MDA (sd-MDA)^2^.

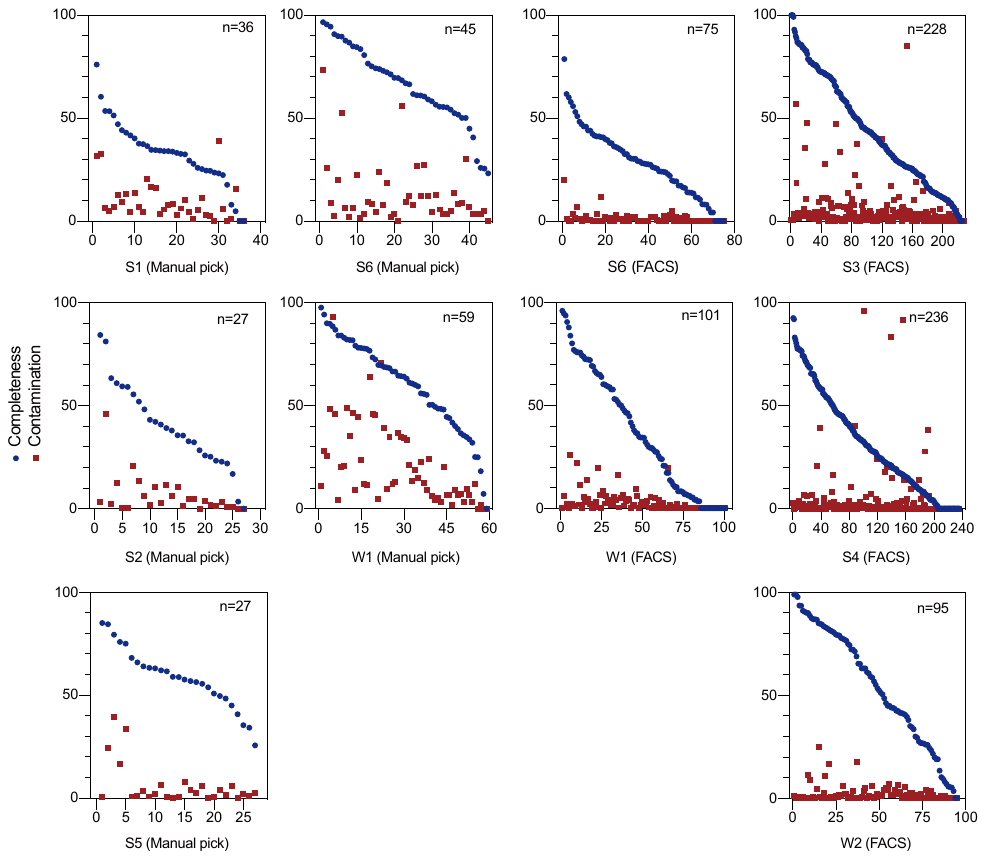

**Supplementary Figure 2. Completeness and contamination statistics for all SAGs obtained from 8 environment samples.**

Genome completeness and contamination of each SAG are plotted by each sampling site. Left boxes are from manual beads picking and right boxes are from FACS-based beads isolation. S6 and W1 were processed both of manual beads picking and FACS-based beads isolation. Samples which were classified as contamination and exhibited 0% completeness were also included.

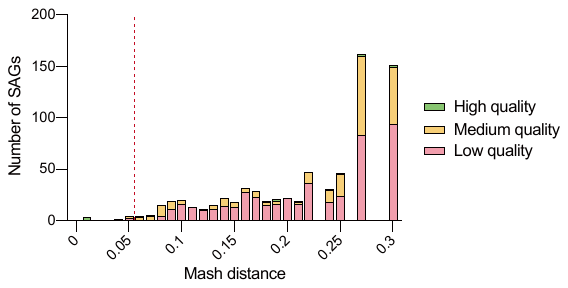

**Supplementary Figure 3. Distribution of mash distance of SAGs obtained from 8 environment samples.**

Mash distance of each SAG to the genomes registered in Refseq was evaluated. The dotted line is the mash distance of 0.05 which can be used as species level threshold.

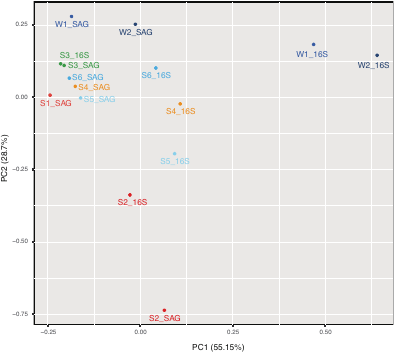

**Supplementary Figure 4. PCA plot for microbial composition in each sampling site and data collection method (16S: 16S rRNA gene sequencing; SAG: SAG-gel).**

Beach (S1) lacks the 16S rRNA gene sequencing data because of the shortage of DNA yield.

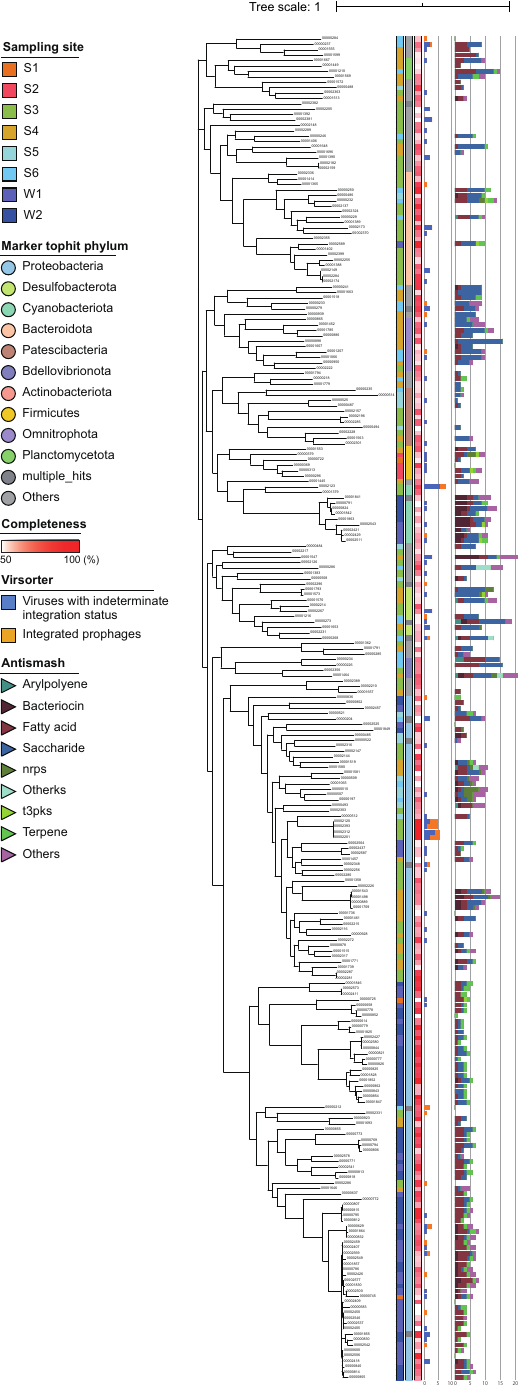

**Supplementary Figure 5. Summary of characteristics of SAGs classified as high-to-medium qualities.**

Genome completeness, taxonomical classification, the number of viral signals and metabolite biosynthetic gene clusters (BGCs) in all sampling sites were summarized. In the result of BGCs, clusters assigned as “cf_putative” were not counted.

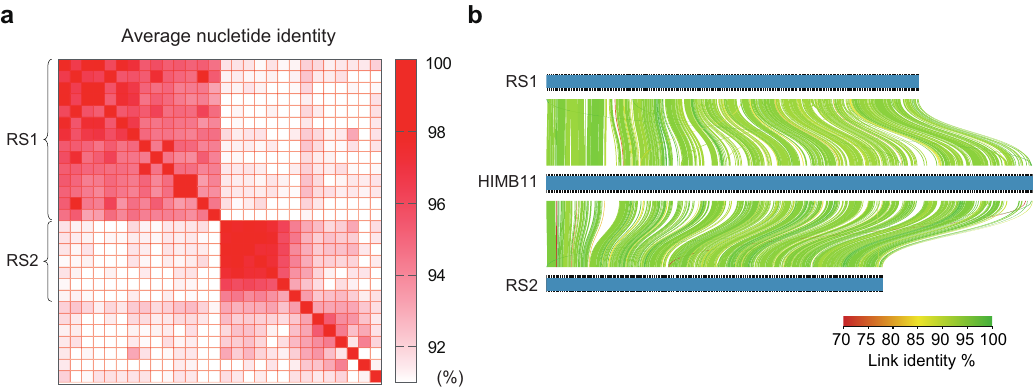

**Supplementary Figure 6. Comparative genome analysis of *Rhodobacter* spp. SAGs obtained from seawater (W1).**

a: ANI of 28 SAGs classified as *Rhodobacter* spp..

b: Sequence alignments were visualized with AliTV.

**Supplementary Table 1. Summary of de novo assembly evaluation with QUAST.**

SAG-gel was compared with conventional in-tube MDA^1^ and previous reported method including compartmented droplet MDA (cd-MDA)^1^, single-droplet MDA (sd-MDA)^2^.

**Supplementary Table 2. Overview of SAG-gel samples**

Number of gel beads processed to 2nd-round amplification, 16S rRNA gene PCR, next-generation sequencing, and quality distributions were summarized for (a)model bacteria and (b)environmental samples.

**Supplementary Table 3. Sequence statistics of 929 sequenced samples**

Summary of sequence statistics including number of contigs, total length, GC%, N50, completeness and contamination, kinds of tRNA, number and length of 5S, 16S, and 23S rRNA gene fragments, taxonomic annotation, and 16S rRNA gene-based Blast search against the Silva database.

**Supplementary Table 4. Summary of mash distance, refseq tophit, and p-value for 929 samples**

**Supplementary Table 5. Summary of enrichment p-value of orthogroups in each sampling site.**

 The cells which exhibited <0.01 p-value are colored by red.

**Supplementary Table 6. List of viral signals.**

**Supplementary Table 7. List of secondary metabolite biosynthetic gene clusters (BGCs).**

**Supplementary Table 8. List of plasmid sequences.**

**Supplementary Table 9. Summary table for the results of genomaple ((a)Complex module, (b)Functional modules, (c)Pathway modules, and (d)Signature modules).**

**Supplementary information**

*SAG-gel with lab-cultured model bacteria*

In order to evaluate the performance of single-cell sequencing, we assessed the SAG-gel with lab-cultured *Escherichia coli* and *Bacillus subtilis*. When we encapsulated *E. coli* at the concentration of 0.1 cell/droplet, the rate of fluorescence-positive beads was 10.5%, suggesting that single cells were successfully reacted with lysis buffer and WGA mixture. In addition, when we generated blank beads that contain no cells, the rate of false-positive beads was less than 0.05%. Hence, we adjusted the cell concentration to 0.1 cell/droplet (3,000 cells/μL) in the following experiment in order to prevent the co-encapsulation of multiple cells in gel beads (0.47% for co-encapsulation). Then gel beads proceeded to FACS-based isolation. With 488 nm laser, a bi-modal fluorescent peak derived from SYBR Green was observed.

After 2nd-round WGA, positive-sorted beads exhibited sufficient DNA amplification for further analysis and sequencing library preparation (> 1.2 μg). The DNA yield was comparable to the manual instruction. In contrast, negative-sorted beads showed no obvious background amplification (< 2.8 ng/μL) (Supplementary Fig. 1a). This indicates that we can easily identify positive and negative samples by endpoint amplicon yield. In our evaluation, > 94% of sorted positive beads showed sufficient DNA amplification for library preparation.

At the result of *de novo* assembling from reads down-sampled to 10× mean mapping depth, the genome cover rate of *E. coli* and *B. subtilis* lysed with enzyme cocktail improved to 51.4% and 64.9%, respectively, while the genome cover rates in conventional alkaline lysis were 42.3% and 39.9%, respectively (Supplementary Fig. 1b). As reported in our previous report, *B. subtilis* showed the superior coverage performance though there were several outliers, which may be attributable to the lower GC content and genome size of *B. subtilis* (43.9%, 4.0 Mbp) compared with those of *E. coli* (50.8%, 4.6 Mbp)^2^. Also, several outliers may be attributed to apoptotic cells, cellular debris, or incompletely lysed cells. The improvement of genome coverage indicated that accessibility of phi29 polymerase to single-cell DNA embedded in the agarose gel matrix and amplification efficiency increased because of the treatment of enzyme cocktail. Especially in *B. subtilis*, the genome cover rate increased by about 25% by enzyme cocktail treatment, suggesting that combining several kinds of lysis method was effective for gram-positive bacteria. In addition, amplification bias was also reduced in enzyme cocktail treatment, which was comparable to conventional in-tube WGA (Supplementary Fig. 1c). The cells embedded in an agarose matrix containing numerous pores, allowing the access of lysis enzymes and detergents while preventing DNA from physical shearing. The washing step can effectively eliminate proteinase, sodium dodecyl sulfate (SDS), and other reagents that inhibit the activity of phi29 polymerase. In addition, because gel matrix is large enough for penetration of enzymes or other components, our SAG-gel-based “reaction and wash” system would be adaptable to other lysis protocols. These characteristics expands the possibilities for improved sequence efficiency of hard-to-lyse bacteria, archaea, and fungi.

The agarose matrix-aided wash and amplification step can minimize the contamination among others between different gel beads. We observed that all of the sequenced SAGs from *E. coli* and *B. subtilis* mixed sample had > 99.6% of their reads mapping to either *E. coli* or *B. subtilis* (Supplementary Fig. 1d). This result outperformed the previously reported single droplet MDA (sd-MDA)^2^, suggesting that contaminating short DNA fragments were removed by washing steps. In addition, the processes from droplet generation to gel bead sorting were conducted with FACS at a normal laboratory environment. This implies that DNA contamination derived from the experimental condition could be prohibited after 1st-round WGA. We also identified the attribution of unmapped reads. They were mainly ascribed to Homo sapiens, Propionibacterium, and Pseudomonas, which are often observed as laboratory contaminants. The number of unmapped reads were equivalent for the results of purified extracted DNA, suggesting that DNA contamination in SAG-gel is not derived from WGA. The *de novo* assembled contigs showed a lower number of misassembled and unaligned contigs in SAG-gel compared to the conventional in-tube MDA because the agarose matrix can capture single-cell DNA while preventing DNA contamination (Supplementary Fig. 1e and Supplementary Table 1). In addition, SAG-gel sequencing has a larger length of N50 and lower number of mismatches and indels, suggesting the accurate genome information can be acquired compared to conventional in-tube MDA.
